# An endosymbiotic *Paenibacillus* sp. modulates disease severity caused by the common watermelon pathogen, *Fusarium oxysporum* f sp. *niveum*

**DOI:** 10.64898/2026.06.18.733246

**Authors:** Dallas Moses, Paola Diaz-Matamoros, Lea Mennen, Lauren Carneal, Kelly Avila, Lina Quesada-Ocampo, Morgan Carter

## Abstract

Fungal plant pathogens can be affected by the bacteria they interact with in their environment, yet the characterization of these interactions beyond direct antagonism is lacking, especially in the case of endohyphal bacteria (EHB). Though limited in characterized examples, EHB can alter disease severity of their fungal host, providing either a potential tool or target for control. We screened isolates of *Fusarium oxysporum* f. sp. *niveum* (FON), an important soil-borne watermelon pathogen, using 16S PCR and fluorescence *in situ* hybridization microscopy to identify novel EHB. A symbiont of FON AS124 was identified to be a *Paenibacillus* sp. through genome sequencing and average nucleotide identity. To begin characterizing this relationship, we conducted watermelon infection assays using FON cured of its symbiont, the native association, and a coinoculation of fungi and bacteria. Disease severity was reduced in watermelon seedlings inoculated with the native association, though not in the coinoculation, and *Paenibacillus* sp. CB74 did not alone promote plant growth or inhibit fungal growth. This study shows an important functional outcome, reduced disease, for a novel symbiosis between FON and *Paenibacillus* sp. CB74, setting up further investigation into the mechanisms behind this outcome and the application of this interaction.

**Importance:** Fungi pose a challenge in both the field and hospital as antifungal resistance rises and chemical control is increasingly scrutinized. In plant pathogenic fungi, endohyphal bacteria may present alternative targets or mechanisms of fungal control. These relationships are observed across diverse groups of fungi and bacteria, though few have been studied to the point of understanding impact. To contribute to the small but growing catalog of known endofungal bacterial relationships, we identified a novel symbiosis and began characterizing its functional outcomes with plant infection assays. The identified bacterial symbiont does alter disease severity of the fungal host offering a new system for both application and study of fungal pathogenesis.

## Introduction

Soil-borne, fungal plant pathogens cause significant loss of crops due to the lack of effective treatments and their ability to persist underground. While factors such as cultivation regimes and moisture levels can influence disease severity in predictable ways, microbial interactions in the rhizosphere can also change plant health outcomes. Bacteria that internally colonize fungal hyphae, endohyphal bacteria (EHB), are reported to increase or decrease virulence, alter host stress responses, and change host production of phytohormones (Richter et al. 2025). While EHB occur across diverse bacterial and fungal groups (Hoffman and Arnold 2010; Robinson et al. 2021), few EHB-host systems have been validated and characterized for functional outcomes. Many fundamental questions about these symbioses remain unanswered, including the range of their impact, though the limited current evidence reveals that EHB can have significant impacts on plant health outcomes when colonizing filamentous plant pathogens (Moses and Carter 2025).

Potentially the most important change EHB induce in hosts is a change in virulence. For example, when the bacterium *Enterobacter* sp. Crenshaw is present in the turfgrass pathogen *Rhizoctonia solani*, disease severity increases (Obasa et al. 2017). Another example is the presence of *Stenotrophomonas maltophilia* within *Fusarium graminearum* increasing the lesion length in wheat seedlings (Ali et al. 2022). While in these two EHB-fungal systems the mechanism driving increased virulence is unknown, in some cases EHB can increase mycotoxin production which is linked to pathogenicity and bacterial-fungal interactions (Venkatesh and Keller 2019). In the case of the cereal pathogen, *F. fujikuroi*, two *Enterobacter* sp. that were identified as EHB increased the expression of the toxin fumonisin (Obasa et al. 2020). However, the increase in fumonisin was not tested *in planta* for changes in disease severity.

*Fusarium* species have arisen as some of the key examples of ascomycete fungi that can harbor EHB, but many agronomically important species complexes have not yet been investigated for bacterial symbionts. *Fusarium oxysporum* poses a challenge across crops, in part because it is a complex of soil-borne filamentous plant pathogens that cannot be easily controlled or treated. For example, the causal agent of Fusarium wilt in watermelon plants, *Fusarium oxysporum* f. sp. *niveum* (FON), is prevalent in the Southeast United States causing watermelon growers yield losses between 30-100% (Amaradasa et al. 2018; Egel et al. 2005). There has been some research into biological agents capable of inhibiting FON, including the identification of a *Bacillus* strain with potential use as a biocontrol (Luo et al. 2023; Xu et al. 2020) and competitive inhibition from a toxin-deficient FON mutant (Xie et al. 2019). Studies focusing on the microbiome interacting with FON have identified similar groups of bacteria co-isolated with the fungal host (Luo et al. 2023) and interacting with FON as core hyphosphere microbiota (Thomas and Antony-Babu 2024). However, the extent and characterization of the interactions between FON and these bacteria have yet to be fully understood to the level needed for biological control outcomes.

In this study, we investigated the hypothesis that FON could harbor internal bacteria that affect the relationship between pathogen and host. We first screened a collection of samples from various watermelon fields and identified a novel endosymbiont that we extracted into pure culture and genome sequenced. Given the interest in the development of new methods for biological control of soil-borne pathogens such as FON, we focused on the impact of this fungal-bacterial interaction on disease severity. We found a reduction in watermelon disease symptoms for the native association between FON and *Paenibacillus* sp. CB74 compared to fungus alone or coinoculation.

## Methods

### Fungal Culturing

FON isolates used in this study were sent to the Carter Lab at UNC Charlotte by the Vegetable Pathology Lab at North Carolina State University. Fungi were stored in water vouchers at room temperature in the dark. FON was cultured at 28 °C on potato dextrose agar (PDA) unless otherwise noted.

### Fungal DNA Extraction and PCR Screening

Fungal isolates were grown in potato dextrose broth (PDB) until mycelium reached between 30mg and 100mg of wet weight. DNA was extracted using the Quick-DNA Fungal/Bacterial Miniprep kit (Zymo) following the protocol with the following change: DNA was eluted using between 50 and 100 µL of molecular grade water. Extractions yielding at least 30 ng/µL of DNA were then tested with PCR to determine if bacterial DNA was present in the sample. PCRs targeting 16S rDNA contained 0.2 µM of forward primer 27F (5’-AGAGTTTGATCTGGCTCAG-3’), 0.2 µM of reverse primer 1492R (5’-GGTTACCTTGTTACGACTT-3’), 2X One Taq Quick Load Master Mix with Standard Buffer (New England Biolabs), and between 200 ng and 400 ng of DNA sample. The samples were run with an annealing temperature of 55°C. Additionally, each isolate was also checked for the quality of DNA extraction by targeting fungal ITS sequences in PCR. Each ITS reaction contained 0.2 µM of forward primer ITS86F (5’-GTGAATCATCGAATCTTTGAA-3’), 0.2 µM of reverse primer ITS4 (5’-TCCTCCGCTTATTGATATGC-3’), 2X One Taq Quick Load Master Mix with Standard Buffer (New England Biolabs), and between 50 ng and 100 ng of DNA sample. The ITS samples were run with an annealing temperature of 53°C. A negative control of no fungal isolate was used through DNA extraction and PCR to check for DNA extraction contamination levels, and a negative PCR control containing molecular grade water in place of DNA template was used to ensure purity of PCR reactions.

### Fluorescent *in situ* Hybridization Screening

Fungal samples grown on water agar slides at room temperature for 3 days were treated with 10% formalin overnight at 4°C. After washing samples with 1X phosphate-buffered saline (PBS) twice, samples were dehydrated using a series of ethanol incubations at 50%, 75%, and 95% for 10 minutes each. Samples were then incubated with 30 µL of hybridization buffer and 1 ngµL^−1^ of Texas Red labeled 16S probe, EUB338 (/TexRed/GCT GCC TCC CGT AGG AGT) at 46°C for 1 hour (Ruiz-Herrera et al. 2015). After washing samples with the hybridization washing buffer twice, Calcofluor white was applied to the samples to visualize the fungal cell wall. After incubating in the dark for about 5 minutes, a Leica Stellaris 8 confocal microscope was used to visualize the samples.

### Bacterial Extraction

To extract bacteria from fungal hyphae, two or three fungal plugs from a 5-7 day old PDA plate were placed in a beaded tube and homogenized for 40 seconds at 20 freq/sec on a TissueLyser II (Qiagen). Samples were filtered through a syringe with a 2.7 µm filter and plated on Lysogeny Broth (LB), King’s Medium B Broth (KB), and Reasoner’s 2 Agar (R2A) plates containing 100 µg/mL cycloheximide. Plates were incubated at 28°C until bacterial growth was observed. Individual colonies were gram-stained.

### Bacterial Genome Sequencing and Assembly

The isolated EHB was grown in KB for 48 hours, and genomic DNA was extracted using the Promega Wizard® Genomic DNA Purification Kit following the gram-negative extraction protocol. Quality of DNA was determined using a NanoDrop Spectrophotometer and the quantity for sequencing using a Qubit Fluorometer (Thermo). Nanopore sequencing and genome assembly was conducted by Plasmidsaurus using v14 library prep chemistry, an R10.4.1 flowcell, and Dorado Super-Accurate Basecalling with default Q10 quality filtering. Default parameters were used for all software except where otherwise noted. The default Plasmidsaurus pipeline was used to assemble the genome which consists of: Filtlong v0.2.1 (github: rrwick/Flitlong) filtering of the bottom 5% fastq reads, downsampling with Miniasm v0.3 (Li 2016) to 100X coverage, Flye v2.9.1 (Kolmogorov et al. 2019) assembly with high quality ONT parameters, and polishing with Mekdaka v1.8.0 (github: nanoproetech/medaka). To annotate the resulting genome, the pipeline beav (Jung et al. 2024) minimal run was used which includes bakta annotation (Schwengers et al. 2021). The (ani_rep) of the Genome Taxonomy Database toolkit (GTDB-Tk) v2.3.2 was run (Chaumeil et al. 2022); FastANI was used in an all-vs-all to visualize average nucleotide identity (Jain et al. 2018). Then the GTDB-Tk *de novo* workflow was run with the bacteria flag, the taxa filter for the genera *Paenibacillus* and *Cohnella*, and the genus *Cohnella* as the outgroup used as options.

### Curing Fungal Isolates of Bacterial Symbionts

To remove *Paenibacillus* sp. CB74 from the FON AS124 isolate for downstream experimentation, FON AS124 was grown on PDA with ciprofloxacin (Cp) at 60 µg/mL for 3 days at 28°C. After 3 days, scrapings from the edge of the fungal growth were used to inoculate a 3 mL PDB culture containing Cp at 60 µg/mL. After 3 days of growth at 28°C in a shaking incubator, the fungal culture was plated on PDB containing Cp at 60 µg/mL. This subculturing was repeated twice for a total of three rounds of culturing on antibiotics. To confirm that the fungus was cured of *Paenibacillus* sp. CB74, fungal mycelia was emulsified and plated on LB with cycloheximide at 100 µg/mL to confirm there were no bacteria present.

### Checking for FON-*Paenibacillus* sp. CB74 inhibition

To test for any direct inhibition of fungal growth by *Paenibacillus* sp. CB74, an agar punch of cured FON AS124 was cultured on 3 PDA plates 2 cm away from 20 µL of a 24-hour LB culture of *Paenibacillus* sp. CB74. Three control PDA plates consisted of only an agar punch of cured FON AS124. Plates were incubated for 4 days at 28 °C.

### Watermelon Infection Trials

To prepare FON inoculum, we grew FON AS124 containing the endosymbiotic *Paenibacillus* p. CB74 and cured FON AS124 on malt extract agar (MEA) plates at 28°C in the dark until the cultures filled the petri dish, usually 5-7 days. To collect conidial suspensions, we added 5 mL of sterile water to each plate, used a pipette tip to aggravate the hyphae, and filtered the suspension through a 40 µm cell strainer. Presence of external *Paenibacillus* sp. CB74 in the conidia suspension of the original association was confirmed with light microscopy. Conidia concentrations were measured using a validated cell counter. Each suspension was adjusted to contain between 4 × 10^6^ and 6 × 10^6^ conidia per mL. For each treatment, 5 mL of conidia suspension was placed in a reservoir. A 48-hour, turbid 12 mL culture of *Paenibacillus* p. CB74 was pelleted and added to a suspension of cured FON AS124 conidia for a coinoculation. The mock treatment consisted of 5 mL of sterile water.

We grew FON susceptible watermelon seedlings (Sugar Baby and Black Diamond, Everwilde Farms, Fallbrook, CA) in Miracle Grow soil to the true leaf stage in a Caron growth chamber at 26°C with 16 hours light daily and 65% humidity. Seedlings were removed from the soil, and their roots were rinsed with DI water until most of the soil was removed. Seedlings were inoculated for 2 minutes by dipping their roots into the mock treatment, cured FON conidia suspension, original FON-*Paenibacillus* sp. CB74 association conidia suspension, or coinoculated with the cured FON conidia suspension and 48-hour *Paenibacillus* sp. CB74 culture then potted individually in Miracle Grow soil and placed in the Caron growth chamber at the same conditions listed above. At two weeks post inoculation, vine lengths of Sugar Baby seedlings were measured from the soil line to the apical node. In a second assay with Black Diamond seedlings, wilt symptoms were rated at the same time point of two weeks using the scale from the NC State Watermelon Disease Handbook to obtain a more holistic measurement of disease progress (Williams and Palmer 1982). Because data from both Sugar Baby vine length measurements and Black Diamond wilt ratings were not normal, Kruskal-Wallis tests were used to determine differences in treatments followed by Dunn’s post-hoc analyses.

To check for impacts on watermelon growth caused by *Paenibacillus* sp. CB74, three trials were conducted using only a 48-hour culture of *Paenibacillus* sp. CB74 on Sugar Baby watermelon seedlings using the same methods described above. Vine length was measured after three weeks and data were analyzed using a Kruskal-Wallis test because the data were not normal.

## Results

### Evidence for endohyphal bacteria in *Fusarium oxysporum* f.sp. *niveum*

Our initial goal was to determine whether bacterial endosymbionts occurred in FON samples from watermelon fields. We screened seven samples isolated during previous studies by the NCSU Vegetable Pathology Lab. PCR screening of DNA prepared from FON isolates revealed three cultures contained bacterial 16S sequences (**Supplementary Table S1**). With FISH microscopy using a universal16S probe, bacterial cells were identified within FON AS124w hyphae (**Figure 1**). Though positive by PCR, both FON 601 and FON AS125 did not exhibit bacteria within or outside of hyphae when visualized with FISH microscopy and were considered inconclusive as to EHB status.

**Figure 1.**
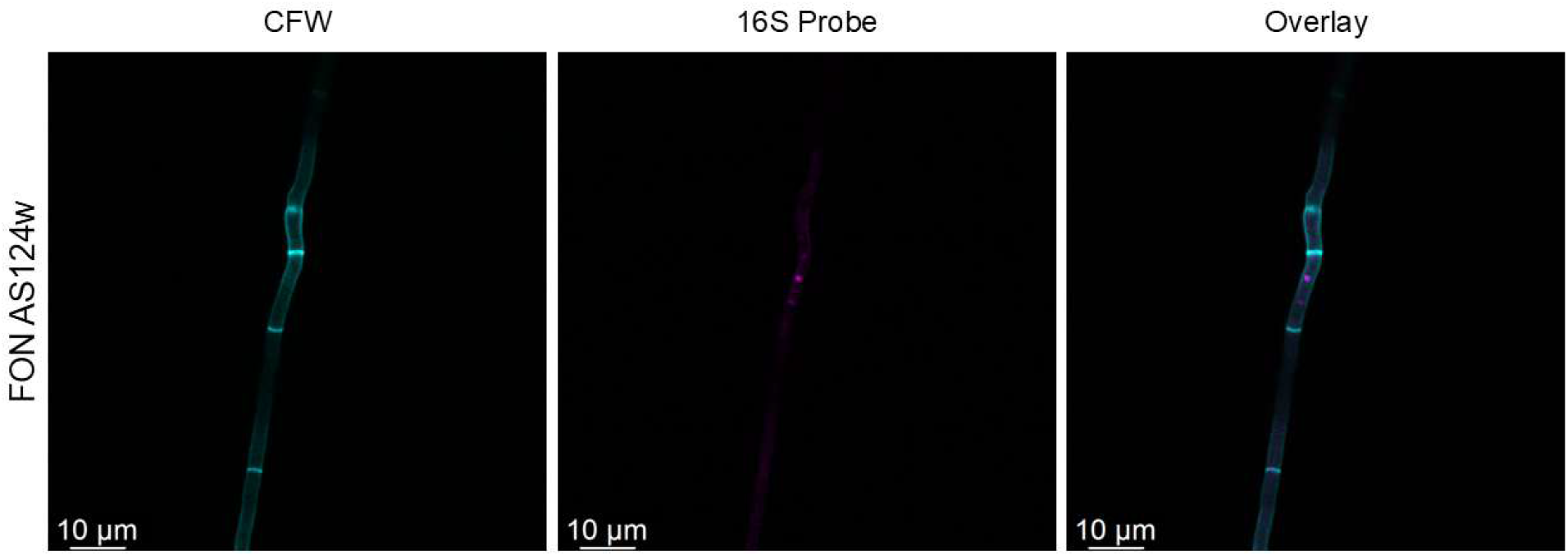
Bacteria identified within *Fusarium oxysporum* f. sp. *niveum* AS124 hyphae. Fluorescence in situ microscopy with a universal 16S probe showing a bacterium (pink) within FON cell walls. CFW, calcofluor white (cyan).

### Endosymbiont is a culturable *Paenibacillus* sp

To identify the endosymbiont from FON AS124, amplicons from the 16S PCR performed on fungal DNA preparations were sequenced and blasted against the NCBI database, revealing the bacterium to be in the *Paenibacillus* genus. To further confirm this identification, we disrupted the fungal tissue and filtered the extract before plating it on various media. Bacterial colonies formed on LB medium and exhibited pale yellow coloring with round, undulate morphology (**Figure 2A**). We consistently observed colonies with the same morphology from three extractions at different times, with no colonies on negative control plates where reagents were used without any fungi to extract from. The bacteria is rod shaped and gram-stained negative, which can occur in the genus *Paenibacillus* despite its taxonomic placement within Bacillota (Shida et al. 1997) (**Figure 2B**). Additionally, no direct inhibition of FON AS124 occurred when plated alongside *Paenibacillus* sp. CB74 (**Figure 2C**).

**Figure 2.**
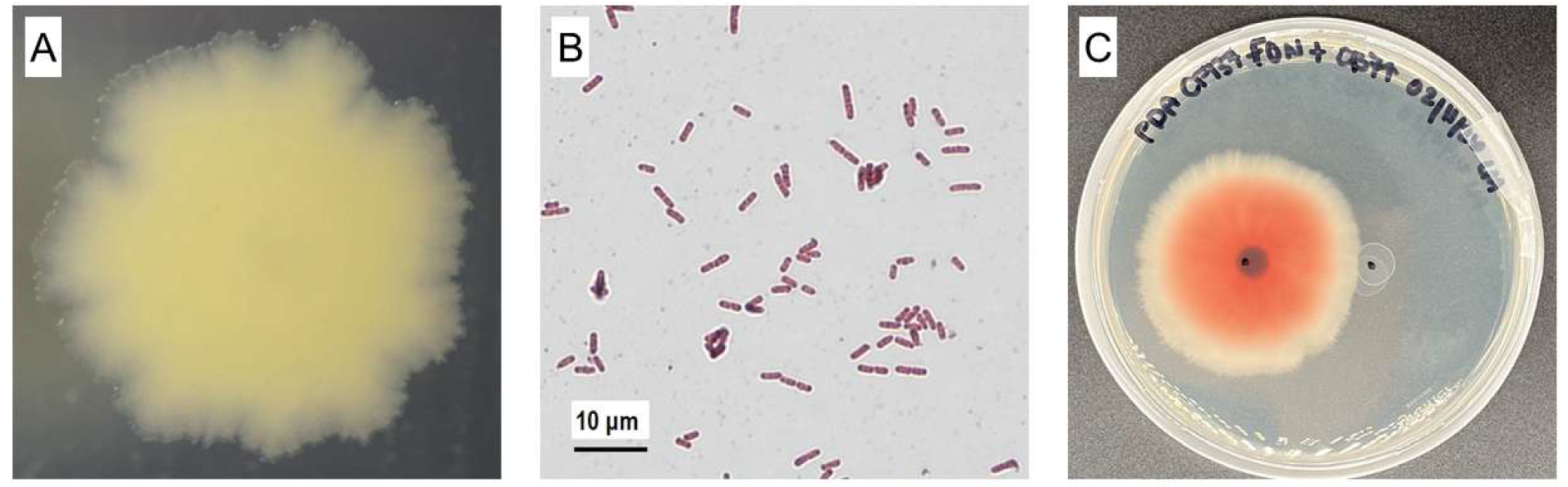
Characterization of the bacterial symbiont from *Fusarium oxysporum* f. sp. *niveum* (FON). (A) Undulate colony morphology of the bacterium extracted from FON on LB agar media. (B) Gram staining of the extracted bacterium revealing gram-negative rod-shaped morphology. (C) Coculture of cured FON AS124 (right) and 20 µL of *Paenibacillus* sp. CB74 (left) on PDA.

### Genomic Analyses

To further characterize the novel endosymbiont, we sequenced the genome with Oxford Nanopore Technologies long-read sequencing obtaining 149,546 raw reads. After processing, our assembly (Accession: JBVMUS000000000.1) with 100X coverage has a circular chromosome of 6,863,592 bp and a second circular replicon of 531,737 bp (**Figure 3A**) with a genomic GC content of 45.6%. When analyzed with the Genome Taxonomy Database toolkit, CB74 falls within the *Paenibacillus* genus, with the most closely related reference strain being *Paenibacillus* sp. 1781tsa1 (GCF_024159265.1) with an average nucleotide identity of 94.5%, just below the typical 95% for species discrimination (**Figure 3B**). Thus, CB74 may represent a new species of *Paenibacillus*, but the taxonomy in this genus is unresolved (**Supplemental Figure S1**).

**Figure 3.**
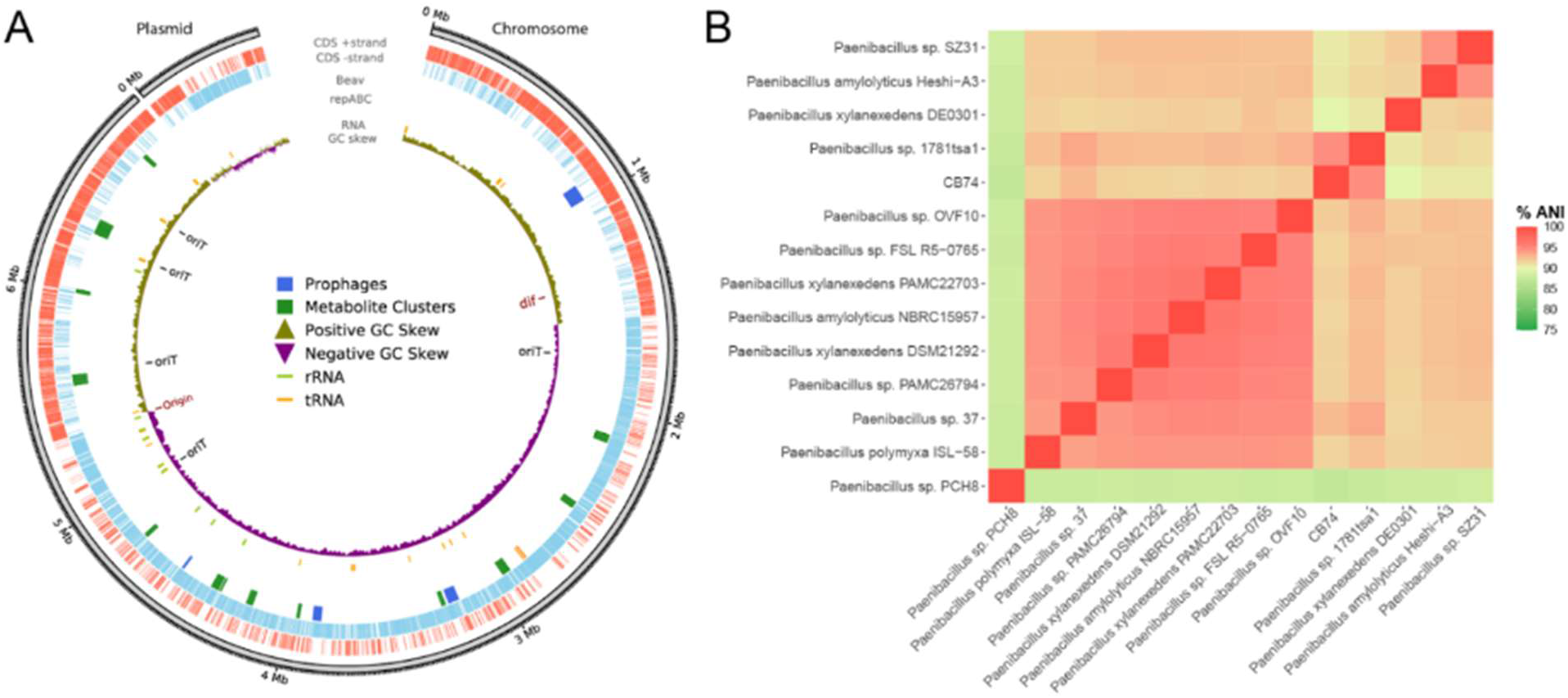
Genomic analyses of the endohyphal *Paenibacillus* sp. CB74. A) Circos plot showing key features of the genome assembly. B) Average nucleotide identity (%ANI) analysis with closely related *Paenibacillus* species as identified by the Genome Taxonomy Database. Typically, 95% is considered the threshold for two strains to be the same species.

The *Paenibacillus* sp. CB74 genome has 5,972 coding sequences, 102 tRNAs, and 36 rRNAs. Additionally, there are eleven putative defense systems and four prophages. The twelve secondary metabolite clusters are all located on the chromosome and are expected to produce two polyketides, two nonribosomal peptides (NRP), a lassopeptide, a proteusin, three metallophores, a terpene, a lanthipeptide, and a NRP hybrid (**Supplementary Table S2**). Though the secondary metabolite clusters have homologs across many *Paenibacillus* species, few are similar to characterized clusters. The two with high known cluster similarity are the zincophore bacillopaline (cluster 12) and the siderophore bacillibactin (cluster 9). It is not apparent whether any of the twelve secondary metabolite gene clusters have direct antifungal activity, though bacillibactin has been correlated with antifungal activity in *Bacillus* spp. (Dimopoulou et al. 2021; Tian et al. 2021). Because some *Paenibacillus* strains have nitrogen fixing capabilities (Xie et al. 2014), we queried the NFixDB to detect nitrogenases or nitrogenase-like genes in CB74 and found none (Bellanger et al. 2024).

### *Paenibacillus* sp. CB74 impacts FON disease severity

The impact of *Paenibacillus* sp. CB74 on FON disease severity was evaluated by inoculating watermelon seedlings with cured FON, FON associated with *Paenibacillus* sp. CB74, and a coinoculation of cured FON and *Paenibacillus* sp. CB74. To rule out any effects on plant growth by *Paenibacillus* sp. CB74, Sugar Baby watermelon seedlings were inoculated with only the bacteria and vine length was measured. Watermelon seedlings inoculated with *Paenibacillus* sp. CB74 did not exhibit any disease symptoms and grew to the same length as the control (p = 0.6345) (**Supplementary Figure S2**). However, differences in disease severity were observed when watermelon seedlings were infected with FON with and without its bacterial symbiont. In Sugar Baby watermelon seedlings, inoculation with the FON-*Paenibacillus* sp. CB74 coinoculum resulted in shorter vine lengths than the mock and FON-*Paenibacillus* sp. CB74 original association, though inoculation with FON did not result in different vine lengths than any of the treatments (p = 0.002056) (**Figure 4A**). Due to the noise observed in vine length and the lack of representation of wilt symptoms from this metric, we repeated the assays with another fully FON-susceptible watermelon variety, Black Diamond. Inoculation with FON and the FON-*Paenibacillus* sp. CB74 coinoculum resulted in higher wilt ratings than the mock and FON-*Paenibacillus* sp. CB74 original association (p = 7.903e-10) (**Figure 4B**). The seedlings inoculated with the original association between FON and *Paenibacillus* sp. CB74 exhibited reduced disease severity, but the coinoculation between FON and *Paenibacillus* sp. CB74 did not result in reduced disease severity compared to cured FON. In both watermelon genotypes, there is a consistent suppression of disease when seedlings are inoculated with the original association between FON and *Paenibacillus* sp. CB74 that is not replicated with the coinoculation.

**Figure 4.**
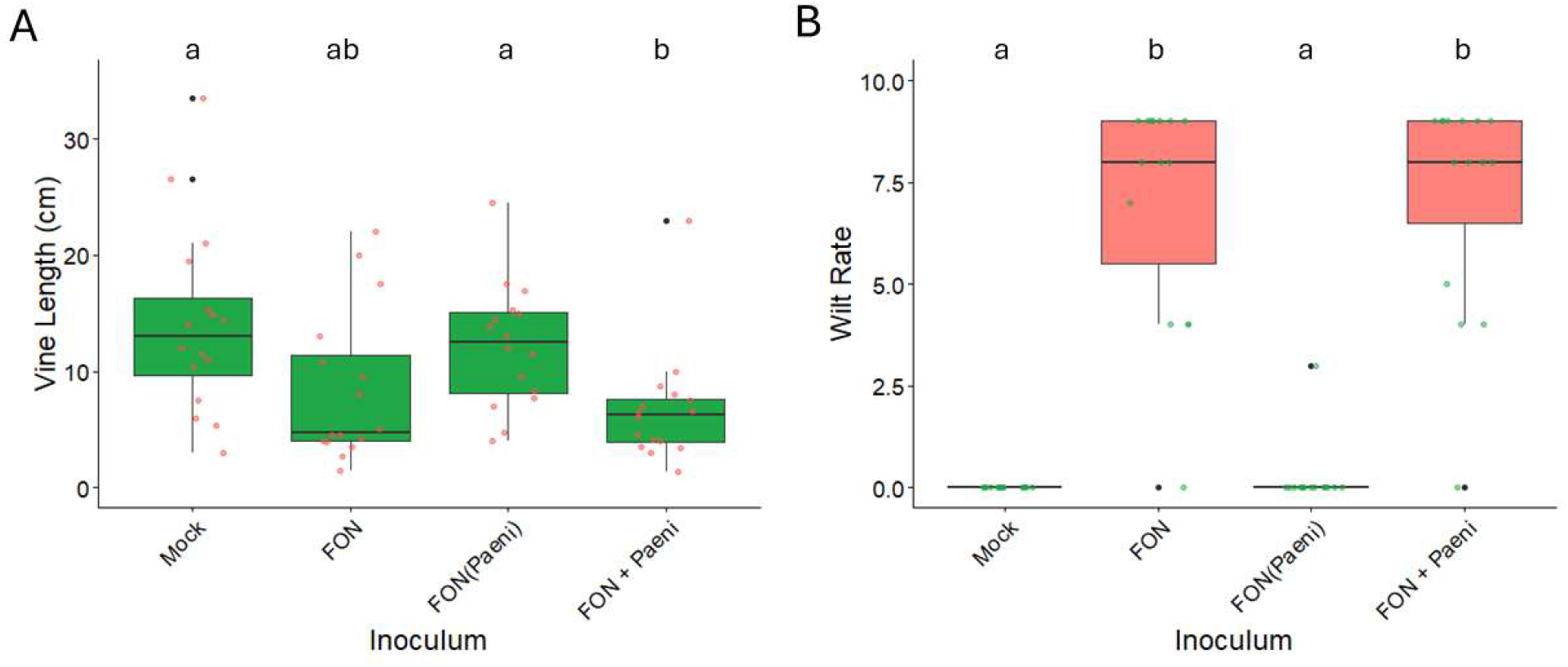
*Fusarium oxysporum* f. sp. *niveum* (FON) disease severity differences in susceptible watermelon varieties. (A) Vine length of Sugar Baby watermelon seedlings at 2 weeks post inoculation. (B) Wilt rating of Black Diamond watermelon seedlings at 2 weeks post inoculation. FON(Paeni) represents a conidial suspension from original association between FON and *Paenibacillus* sp. CB74. FON + Paeni represents cured FON conidia combined with a suspension of *Paenibacillus* sp. CB74. Significant differences in vine length and wilt rating are noted by lowercase letters and were determined with Kruskal-Wallis rank sum tests and Dunn’s post-hoc tests.

## Discussion

We identified a novel EHB, *Paenibacillus* sp. CB74, that decreases disease severity caused by FON. Though no direct antagonism was observed between the bacterium and fungus, the original isolate in long-term symbiosis with *Paenibacillus* sp. CB74 reduced wilt symptoms and increased vine length in watermelon varieties fully susceptible to FON. The outcomes of this EHB-fungal relationship emphasize the importance of understanding the complex microbe-microbe interactions that impact plant-microbe interactions, particularly plant-pathogen interactions. Fusarium spp. like FON are difficult to control in the field but investigating the microorganisms that exist around and inside of them has already uncovered new avenues of potential control. With this relatively small-scale screening of FON, we were able to identify a bacterium with a negative impact on virulence outcomes that has the potential for biocontrol applications in this pathosystem and possibly beyond.

Our discovery is in line with other studies that have found important outcomes of bacterial-Fusarium interactions but highlights that there is much diversity that is still unexplored (Luo et al. 2023; Obasa et al. 2020; Thomas and Antony-Babu 2024; Xu et al. 2020). In particular, the FON-*Paenibacillus* sp. CB74 microbial relationship results in reduced disease severity which is different than other documented cases of EHB affecting virulence positively (Ali et al. 2022; Obasa et al. 2020; Obasa et al. 2017). Additionally, reduced disease in watermelon could not be replicated by simply coinoculating with both FON and *Paenibacillus* sp. CB74. In contrast, the reestablishment of the fungus-EHB relationship after curing the fungal host restored increased virulence in the *F. fujikuroi* and *Enterobacter* sp. relationship (Obasa et al. 2020). Researchers were able to link increased fumonisin production with the presence of the EHB partner which is thought to be responsible for increased virulence. Investigating the FON genome and identifying differences in gene expression while in symbiosis with *Paenibacillus* sp. CB74 may yield potential hypotheses explaining the observed differences in disease severity.

The genus *Paenibacillus* has long been known for being common in the rhizosphere and phytobiome (McSpadden Gardener 2004; Rybakova et al. 2016). Academic and commercial interest in the genus have discovered strains with the potential for plant growth promotion, nitrogen fixation, and antimicrobial activity (Dobrzynski and Nazieblo 2024; Grady et al. 2016; McSpadden Gardener 2004). Our investigation of *Paenibacillus* sp. CB74 does not ascribe these qualities to this strain, though there are limitations in our experiments that may have obscured them. Inoculation with *Paenibacillus* sp. CB74 alone did not promote watermelon seedling growth in a growth chamber. Supporting that observation, the strain does not have putative nitrogenases or genes associated with nitrogen fixation. Finally, we did not see a direct antagonism between *Paenibacillus* sp. CB74 and FON, though it may have antifungal or antibacterial activity against other untested microbes. The finding that *Paenibacillus* sp. CB74 had colonized FON indicates a more nuanced relationship than direct chemical antagonism that is most popularly pursued in biocontrol candidates (Dobrzynski and Nazieblo 2024; Lorentz et al. 2006). That the secondary metabolite repertoire consists of largely uncharacterized compounds leaves open the possibility of antimicrobial or plant interactions that were not captured in our study.

Part of the motivation for conducting this work was to develop new study systems to probe the unknowns of intimate bacterial-fungal interactions within the phytobiome (Moses and Carter 2025). One observed phenomenon is that some bacteria change cell shape within fungal hyphae, as was the case in our observations here where the rod-shaped *Paenibacillus* sp. CB74 appeared more circular in FISH imaging. Additional work is needed to investigate how and whether the bacteria are metabolically active within fungi, and how stresses within the fungal cell may be contributing to the changes in bacterial cell shape. Another outstanding question is that of host specificity and whether *Paenibacillus* sp. CB74 can colonize other FON, *Fusarium*, or fungi more broadly. As this is a facultative interaction, we hypothesize that *Paenibacillus* sp. CB74 will be able to colonize other fungi, such as what is observed with the EHB *Luteibacter mycovicinus* and its colonization of Ascomycota hosts (Arendt et al. 2016). The application of reassociation assays where microbes are cocultured under nutrient stress will likely yield additional insights into symbiosis formation and specificity (Arendt et al. 2016).

Ultimately, our study adds to the limited yet growing set of findings that underscore the importance of intermicrobial interactions on plant health. In particular, the often-overlooked phenomenon of bacteria colonizing fungal hyphae intracellularly can create disease outcomes that are potentially beneficial for agricultural systems. Beyond the development of biocontrols, we also view interactions like *Paenibacillus-*FON as a key to unraveling mysteries of fungal biology. Further characterization of the mechanisms of disease suppression may reveal more insights into the basics of pathogenicity and fungal cell biology. As we discover more of these relationships, we can begin to identify the rules and conditions resulting in endohyphal establishment and disease control.

## Acknowledgements

Support was provided by the USDA National Institute of Food and Agriculture (project award no. 2021-67034-35134) and the North Carolina Biotechnology Center (no. 2026-FLG-0206) to MEC.

## Data Availability

This Whole Genome Shotgun project (PRJNA1402388) has been deposited at DDBJ/ENA/GenBank under the accession JBVMUS000000000. The version described in this paper is version JBVMUS010000000.

